# High-Resolution Melting (HRM) Assay for Molecular Detection of *Calonectria ilicicola* in Soybean

**DOI:** 10.64898/2026.09.01.748390

**Authors:** Krishna R. Pandey, Boris X. Camiletti, Jeannie M. Klein-Gordon, Nathan E. Schroeder, Steven J. Clough, Hari S. Karki

## Abstract

*Calonectria ilicicola*, an emerging and economically important soilborne pathogen in the U.S. Midwest, causes red crown rot (RCR) disease of soybean. It is essential to have an early and accurate detection method for effective management of RCR, since disease symptoms are nearly identical to those of other soybean diseases. In this study, we developed and validated a high-resolution melting (HRM) assay targeting the translation elongation factor 1α (TEF-1α) gene for the specific detection of *C. ilicicola*. The assay was evaluated using a specificity panel of 78 DNA samples, including 67 fungal and oomycete DNA samples and 11 soybean host-DNA samples, as well as DNA from 56 field-collected symptomatic and asymptomatic soybean tissues. The HRM assay generated distinct melting profiles and, where applicable, absence or delayed amplification that reliably differentiated *C. ilicicola* from non-target fungi and oomycetes. Additionally, the marker distinguished three haplotypes among *C. ilicicola* isolates, consistent with nucleotide polymorphisms in the TEF-1α target region. The assay demonstrated strong quantitative performance, exhibiting high linearity (R² = 0.9992) and acceptable amplification efficiency (94.05%) across a broad DNA concentration range. In field samples, HRM-based detection aligned with disease status, accurately identifying *C. ilicicola* in symptomatic plants and no detection in asymptomatic tissues. Overall, this HRM assay provides a rapid, sensitive, and reliable diagnostic tool for routine molecular detection of *C. ilicicola*.

## Introduction

*Calonectria ilicicola* is a soilborne fungal pathogen that causes red crown rot (RCR) of soybean and Cylindrocladium black rot of peanut. In soybean, RCR has become an increasing concern in the major U.S. production regions, including the Midwestern states, where the disease has been reported in Illinois, Kentucky, Indiana, Missouri, Minnesota and Wisconsin (Bish et al. 2025; Bonkowski et al. 2024; Clough et al. 2026; Groves et al. 2026; Kleczewski et al. 2019; Malvick et al. 2026; Neves et al. 2023). Field diagnosis of RCR can be challenging because its symptoms often overlap with those caused by other soybean diseases and abiotic stresses. Although reddish discoloration of the crown region and the presence of red perithecia are good diagnostic indicators of RCR, these features are not always visible under field conditions (Telenko and Bonkowski 2024). Foliar symptoms, including chlorosis and interveinal necrosis, resemble those caused by sudden death syndrome (SDS), brown stem rot, and southern stem canker (Kleczewski et al. 2022). Therefore, visual diagnosis alone may not reliably identify RCR in the field. Diagnosis is further complicated by mixed infections, as a prior study identified that most soybean root samples (86%) contained two to six pathogen species, while only 11% carried a single species, with frequent co-occurrence of *Fusarium virguliforme* (causal agent of SDS) and *C. ilicicola* in the same diseased tissues (Ye et al. 2020). Traditional diagnostic methods typically involve examining symptomatic tissues, isolating the suspected pathogen on culture media, and identifying the organism using morphological characteristics and molecular approaches such as PCR and Sanger sequencing of a few loci (Parkinson et al. 2019). However, morphology-based identification can be challenging for closely related fungi (Parkinson et al. 2019; Schoch et al. 1999) and molecular workflows that require isolation, subculturing, DNA extraction, PCR, gel electrophoresis, and sequencing may take several days or longer (Niessen 2015; Notomi et al. 2015). Therefore, faster and more sensitive molecular approaches are needed for early and accurate disease detection (Fang and Ramasamy 2015; Sankaran et al. 2010).

Few DNA-based assays have been developed to detect *C. ilicicola*. Loop-mediated isothermal amplification (LAMP) assays targeting the β-tubulin gene have been designed for detection of *C. ilicicola* in avocado and soybean (Lu et al. 2022; Parkinson et al. 2019). In addition, a quantitative PCR (qPCR) assay based on the intergenic spacer (IGS) region of ribosomal DNA was developed to detect and quantify *C. ilicicola* in naturally infested soil samples (Ochi and Kuroda 2021). These studies show that molecular tools can provide sensitive and specific detection of *C. ilicicola*. However, some of these approaches require post-amplification processing, sequencing, specialized probes, or involve risks such as non-specific amplification, challenges in assay design, and additional costs, which can limit their practicality when large numbers of samples must be processed (Crego-Vicente et al. 2024; Yang et al. 2024).

High-resolution melting (HRM) is a rapid method that rapidly distinguishes DNA fragments based on sequence variation using a closed-tube format. HRM works by monitoring the melting behavior of double-stranded DNA as temperature increases, causing a fluorescence decrease as the strands separate (Montgomery et al. 2007). Even minor sequence differences generate distinct melting profiles, making HRM a sensitive approach for detecting genetic variation (Liew et al. 2004; Reed and Wittwer 2004). In addition, HRM does not require labeled probes or post-PCR processing, making it useful for high-throughput diagnostic applications where rapid pathogen differentiation is needed (Muneeswaran et al. 2024; Sahoo et al. 2025). HRM has been widely applied in plant pathology for rapid pathogen detection, species discrimination, and genotyping. For example, HRM has been used to differentiate closely related *Fusarium* species causing ear rot in maize (Schiwek et al. 2020), to enable species-level identification of *Phytophthora* (Zambounis et al. 2016), to detect *Alternaria* associated with citrus brown spot (Garganese et al. 2018), and to resolve population-level genetic differences in *Venturia inaequalis* (Chatzidimopoulos et al. 2019). Additional applications include distinguishing *Phytophthora sojae* avirulence alleles and differentiating *Fusarium graminearum* chemotypes, demonstrating HRM’s versatility for characterizing plant-pathogenic organisms (Santhanam et al. 2024; Singh et al. 2024).

Selection of an appropriate genetic marker is critical for species-level discrimination of plant pathogens. Although the internal transcribed spacer (ITS) region is widely used as a universal fungal barcode (Blaalid et al. 2013; Schoch et al. 2012), it does not always provide sufficient resolution among closely related species. In contrast, the translation elongation factor 1-alpha (TEF-1α) gene has been widely used in fungal systematics and phylogenetic studies because it offers greater discriminatory power (O’Donnell et al. 1998; Wulff et al. 2010) and is well represented in public reference databases (Henry et al. 2022; O’Donnell et al. 2009). Therefore, TEF-1α was selected as a possible target region for developing an HRM-based assay to detect *C. ilicicola* from soybean samples. In this study the TEF-1α sequence was used to develop and validate an HRM-based assay capable of detecting *C. ilicicola* directly from soybean tissues exhibiting root and crown rot or RCR-like symptoms, without the need for pathogen isolation and culturing.

## Materials and Methods

### *C*. *ilicicola* isolates and culture conditions

The Pike County, Illinois *C. ilicicola* isolate CC1/NRRL 64312 (Kleczewski et al. 2019) was used in this study as the reference isolate. Long-term storage of the isolate was maintained on infected barley grains as previously described (Ochi and Nakagawa 2010). For experimental assays, CC1 and other *C. ilicicola* isolates were grown on potato dextrose agar (PDA) plates at room temperature (21-23°C) for 7 days and used for subsequent assays. All fungal and oomycete isolates included in this study are listed in Supplementary Table 1. Non-*C*. *ilicicola* Isolates were grown on PDA plates at room temperature for 10–14 days to obtain sufficient mycelial growth for downstream DNA extraction.

### DNA extraction from fungal, oomycete isolates and infected soybean tissues

Genomic DNA from fungal and oomycete cultures, as well as from infected soybean tissues, was extracted using the Quick-DNA Fungal/Bacterial Miniprep Kit (Zymo Research, USA) following the manufacturer’s instructions. For fungal and oomycete cultures, mycelia were harvested from PDA plates overlaid with sterile cellophane to avoid agar contamination. For field-collected soybean tissues, approximately 50 mg of scraped crown tissue was dried overnight in a SpeedVac concentrator. The dried tissue was homogenized with two sterile 3.2-mm steel beads using a BeadBlaster™ 24 Microtube Homogenizer (Benchmark Scientific, USA) at speed setting 5 for 1 min 30 s. After homogenization, 750 µL of the BashingBead™ buffer was added, and samples were homogenized again under the same settings for 1 min 30 s. The resulting lysates were then processed for DNA extraction according to the kit protocol. Extracted genomic DNA was quantified using a NanoDrop spectrophotometer and adjusted to approximately 1 ng/µL for pure-culture DNA samples and 20 ng/µL for infected soybean tissue DNA samples, unless stated otherwise.

## Primer design

We considered two key biological scenarios when selecting a target region and designing primers for our HRM assay. First, primer-binding sites in non-target fungal species may contain sufficient sequence variation to produce mismatches, resulting in failed or delayed amplification. Second, primer-binding sites may be conserved among *C. ilicicola* isolates, while single-nucleotide polymorphisms (SNPs) within the amplified region could generate distinct HRM profiles, allowing isolate differentiation. To accommodate both scenarios, we evaluated multiple loci previously used for *Calonectria* identification and phylogenetic analyses, including actin (*act*), calmodulin (*cmdA*), histone H3 (*his3*), intergenic spacer (IGS), internal transcribed spacer (ITS), translation elongation factor 1-alpha (TEF-1α), and β-tubulin (*tub2*) (Clough et al. 2026; Liu et al. 2020; Lombard et al. 2010; Ochi and Kuroda 2021). Following sequence analysis of these loci, we identified TEF-1α as the region that best satisfied both primer-design scenarios. The TEF-1α sequences for common soybean root pathogens (*Pythium*, *Phytophthora*, *Fusarium*), as well as *Calonectria* spp. and *C. ilicicola*, were retrieved from NCBI and aligned in Geneious Prime Version 2025.2.1 to assess sequence diversity and conservation across common soilborne pathogens prevalent in soybean fields. To further validate the target region, we sequenced the TEF-1α locus from representative *Calonectria* species and multiple *C. ilicicola* isolates. Based on these sequences, a 3′ portion of TEF-1α was selected, and primers (TEF1-HRM_F: 5′-GTA CGA TGT CAC CGT CAT TGG TAA G-3′ and TEF1-HRM_R: 5′-TTG AGA GTC GGT TAG AGG CTG AAG-3′) were designed in Geneious Prime to amplify a 73-bp fragment.

For primer design in the conventional PCR assay, we targeted the IGS region. However, because only a limited number of IGS sequences for *Calonectria* species are available in NCBI, additional sequences were needed for sufficient analyses. Using the primers described by Ochi and Kuroda (2021), the IGS region was amplified and sequenced from the *Calonectria* species listed in Supplementary Table 1. The resulting PCR products were sequenced by WideSeq and assembled at the Purdue Genomics Center. Sequences were mapped and aligned in Geneious Prime, where a genetic region unique to *C. ilicicola* was identified. New IGS-specific primers (IGS-DNA_F: 5′-GAA GAA GAA CTT GCC CG TC-3′ and IGS-DNA_R: 5′-CTT CTT CCT GCC ACC AAG TA-3′) were then designed in Geneious Prime to amplify a 555-bp fragment.

## HRM and PCR assay conditions

A LightCycler 480 II (Roche, Basel, Switzerland) equipped with a 384-well reaction module was used to carry out all HRM assays. Each 10 µL reaction contained 5 µL of 2× EvaGreen qPCR Master Mix (Biotium, Fremont, CA), 0.5 µM of each primer, 1 µL of genomic DNA template, and nuclease-free water to bring the reaction to the final volume. Cycling conditions followed the LightCycler 480 SYBR Green protocol with minor modifications: an initial denaturation at 95°C for 5 min, followed by 45 cycles of 95°C for 10 s, 61°C for 10 s, and 72°C for 10 s. Melting curve analysis was conducted immediately after amplification. The melt program consisted of 95°C for 5 s, cooling to 65°C for 1 min, and a gradual increase to 97°C with continuous fluorescence acquisition, followed by final cooling at 40°C. Positive and negative controls were included in every run to ensure accurate sample classification, assess assay specificity, and confirm the absence of contamination. All samples were analyzed using at least four technical replicates, and each experiment was repeated twice.

In the conventional PCR assay, reactions were prepared using GoTaq® Green Master Mix (Promega, WI, USA) in a total volume of 25 µL, consisting of 12.5 µL of 2× GoTaq® Green Master Mix, 1 µL of template DNA, 0.5 µM of each primer, and nuclease-free water to bring the reaction to the final volume. Amplification was performed on a Bio-Rad T100™ Thermal Cycler (Bio-Rad Laboratories, Hercules, CA, USA) with an initial denaturation at 95°C for 2 min, followed by 35 cycles of 95°C for 15 s, 56°C for 15 s, and 72°C for 30 s, and a final extension at 72°C for 5 min. PCR products were resolved on 1% agarose gels prepared in 1× TAE buffer, stained with SYBR® Safe DNA Gel Stain (Thermo Fisher Scientific, Waltham, MA, USA), and visualized under blue-light illumination using a Bio-Rad GelDoc Go imaging system. Amplicons producing bands of the expected size were purified using a QIAquick PCR Purification Kit (Qiagen, Hilden, Germany) prior to Sanger sequencing.

## Sensitivity and standard curve analysis

To determine the sensitivity of the HRM assay using primers targeting the TEF-1α gene, a ten-fold serial dilution series of *C. ilicicola* genomic DNA was prepared, ranging from 70 ng/µL to 0.00007 ng/µL. Each dilution was replicated with four technical replicates under the HRM conditions described previously. Cp values were measured for each reaction and used to generate a standard curve by plotting Cp values against the logarithm of DNA concentration. Detection across the dilution series was evaluated based on reproducible amplification and the presence of a consistent and specific HRM melting profile. Amplification efficiency, also referred to as PCR efficiency, was calculated using the equation E = (10^−1/slope^ − 1) × 100%, as recommended in the MIQE guidelines (Bustin et al. 2009). Primer efficiency in the standard-curve analysis was evaluated using widely accepted qPCR quality criteria, including a slope between – 3.1 and –3.6 (corresponding to 90–110% efficiency) and strong linearity indicated by an R^2^ value close to 0.99 (Bustin et al. 2009; Raymaekers et al. 2009).

## HRM assay specificity evaluation

Specificity of the primer set was assessed using a panel of 78 DNA samples, which included 20 *C. ilicicola* isolates, 14 DNA samples from other *Calonectria* species, 33 DNA samples from other fungal and oomycete organisms predominantly associated with soybean or soilborne disease systems, and 11 soybean host-DNA samples representing different cultivars (Supplementary Table 1). A *C. ilicicola* positive control and a no-template water control were included in each run. For the pure-culture specificity panel, samples were designated as positive when they produced reproducible amplification within 30 cycles and exhibited HRM profiles consistent with the *C. ilicicola* positive control. Amplification signals appearing after 30 cycles were interpreted as late or non-specific amplification and were not considered indicative of true target detection unless supported by a reproducible *C. ilicicola*-like HRM profile.

### HRM data processing and visualization

Initial visualization of amplification curves, melting curves, derivative melt peaks, melting temperatures, and Cp values was performed using LightCycler 480 II software. HRM profiles were further examined using uAnalyze (Dwight et al. 2012) to compare melting curve patterns and peak separation among target and non-target samples. For figure preparation, raw fluorescence data and derivative melt data were exported from the LightCycler 480 II software and replotted in R version 4.5.1 to generate amplification plots, melting curves, and derivative melt peak plots. Standard curve analysis was also performed in R version 4.5.1 using Cp values obtained from the serial dilution assay, and amplification efficiency was calculated as described in the previous section. Summary metrics, including mean melting temperature and standard deviation, were computed from technical replicates in R version 4.5.1.

## Results

### Sequence variation in selected loci validates *C. ilicicola*-specific assay development

Sequence analysis of the TEF-1α gene from major soybean soilborne pathogens (*Fusarium*, *Pythium*, *Phytophthora*), multiple *Calonectria* species, numerous *C. ilicicola* isolates retrieved from the NCBI nucleotide database, publicly available whole-genome shotgun (WGS) assemblies in the NCBI, and in-house TEF-1α gene sequencing demonstrated that the HRM primer-binding sites were highly conserved within all *C. ilicicola* isolates but exhibited substantial nucleotide divergence in non-target fungi and oomycetes. Within the resulting 73-bp TEF-1α amplicon, three haplotypes were detected among *C. ilicicola* isolates. In-house sequencing showed that isolates from Illinois and surrounding states consistently possessed a guanine SNP (Type I), whereas a small number of historical and one Hawaiian isolate carried an adenine at the same position (Type III). Three Hawaiian isolates contained a single-nucleotide deletion at the reverse-primer binding site (Type II) while retaining the Type I SNP. Representatives sequences for these Hawaiian isolates have been deposited in NCBI under accession numbers PZ834570 and PZ834571. With our search in the NCBI nucleotide sequences of TEF-1α, only two haplotypes were available in which 63 were Type I and 25 were Type III based on a random selection of 88 TEF-1α nucleotide sequences and, across all 38 publicly available *C*. *ilicicola* whole-genome shotgun (WGS) assemblies, 21 isolates represent Type I and 17 isolates represent Type III sequences (Supplementary Table 2). Melt-curve simulations using uMELT (Dwight et al. 2011) predicted distinct HRM profiles among these variants, further supporting the capacity of the TEF-1α amplicon to resolve both species identity and isolate-level haplotypes. In parallel, analysis of the 2.3-kb intergenic spacer (IGS) region across multiple *Calonectria* species identified an IGS sequence unique to *C. ilicicola*. IGS sequences generated in this study were deposited in NCBI GenBank under accession numbers PZ824507–PZ824517. Alignment of newly generated and publicly available IGS sequences revealed consistent polymorphisms that clearly differentiated *C. ilicicola* from closely related taxa. Leveraging this region, a conventional PCR primer pair was designed that amplified a 555-bp *C. ilicicola*-specific fragment, providing an additional assay for species confirmation through amplicon size and sequence analysis.

## Standard curve and sensitivity of the HRM Assay

To assess the analytical sensitivity of the HRM assay, a standard curve was generated from serial dilutions of *C. ilicicola* genomic DNA ranging from 70 ng/µL to 0.00007 ng/µL. Mean crossing point (Cp) values from four technical replicates were plotted against the log10 of DNA concentration (Supplementary Figure 2). The standard curve exhibited strong linearity across the dilution range, with a regression equation of y = −3.473x + 21.1, a coefficient of determination (R² = 0.9992), and a slope-derived amplification efficiency of 94.05%. Cp values increased consistently with decreasing DNA concentration, ranging from 14.8 cycles at 70 ng/µL to 36 cycles at 0.00007 ng/µL. At the assay cutoff of 30 cycles, the HRM assay reliably detected DNA concentrations as low as 0.007 ng/µL. Amplification curves for all dilutions showed clear concentration-dependent separation (Supplementary Figure 2), supporting the linear relationship observed in the standard curve and confirming reliable detection across a broad dynamic range.

## Differentiation of *C. ilicicola* from non-target fungi, oomycetes, and soybean DNA by HRM

To evaluate assay specificity, 78 DNA samples were analyzed, including 67 fungal and oomycete DNA samples and 11 soybean host-DNA samples representing different cultivars (Supplementary Table 1). All *C. ilicicola* DNA samples produced robust amplification with HRM profiles consistent with the *C. ilicicola* positive control and Cp values clustering ∼20 cycles (Figure 2). A small subset amplified later around 25-27 cycles but still displayed the characteristic *C. ilicicola* HRM profiles. Across the entire sample set, amplification within the ∼30-cycle threshold occurred only for *C. ilicicola* isolates and a single non-target species, *C. chinensis*; however, *C. chinensis* displayed a distinctly different melting peak and curve compared with *C. ilicicola* (Figure 2). All other non-target fungi and oomycetes, including *Fusarium*, *Pythium*, and *Phytophthora*, either failed to amplify or produced only late, inconsistent amplification beyond the diagnostic range (Supplementary Figure 3). Soybean host-DNA samples showed delayed amplification (>35 cycles) with non-specific and irreproducible melt peaks, and the no-template water control showed no amplification. Together, these results demonstrate that the HRM assay distinguished *C. ilicicola* from the non-target fungi, oomycetes, and soybean host-DNA samples tested in this study.

**Figure 1.**
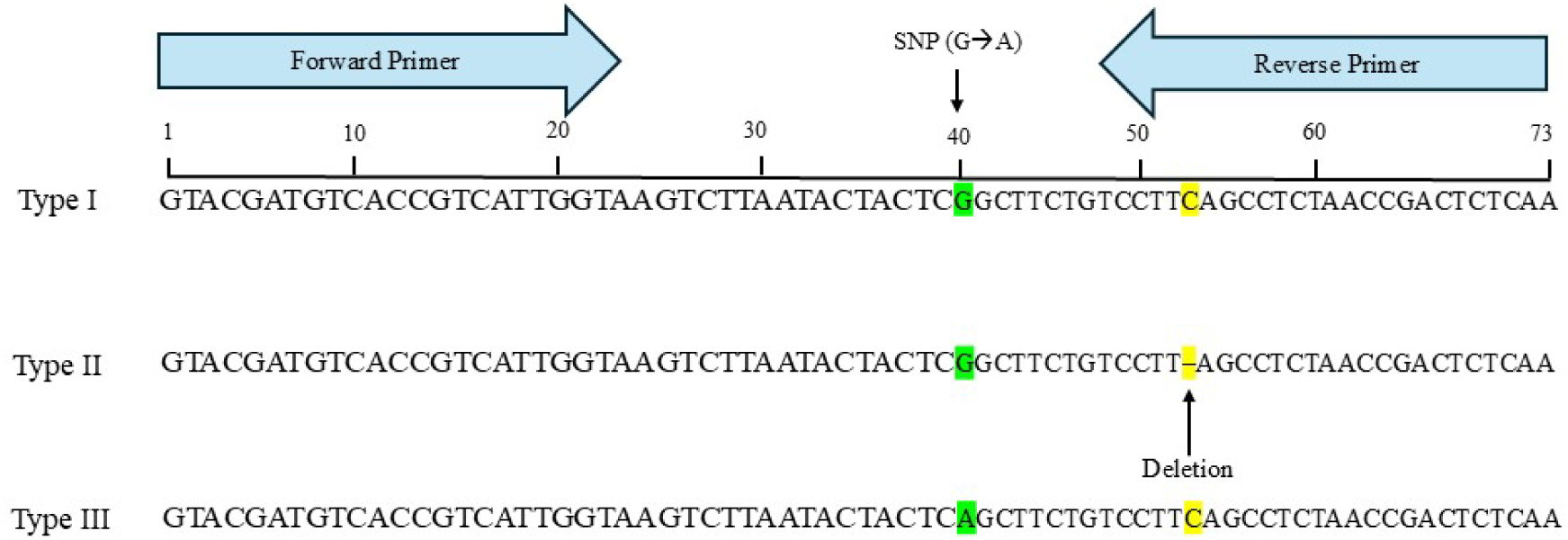
Targeted TEF-1α gene region used for HRM primer design for *C. ilicicola*. The aligned nucleotide sequences represent three sequence variants (Type I, Type II, and Type III) within the TEF-1α gene of *C. ilicicola*. Variants differ by a SNP (G→A) and a nucleotide deletion within the reverse primer binding site. Forward and reverse HRM primers were positioned to flank this SNP segment to maximize melt-curve resolution and enable discrimination among the isolates. Numbered positions (1–73) correspond to nucleotide coordinates within the amplified fragment. These polymorphisms form the basis for differentiating *C. ilicicola* isolates using HRM analysis.

**Figure 2.**
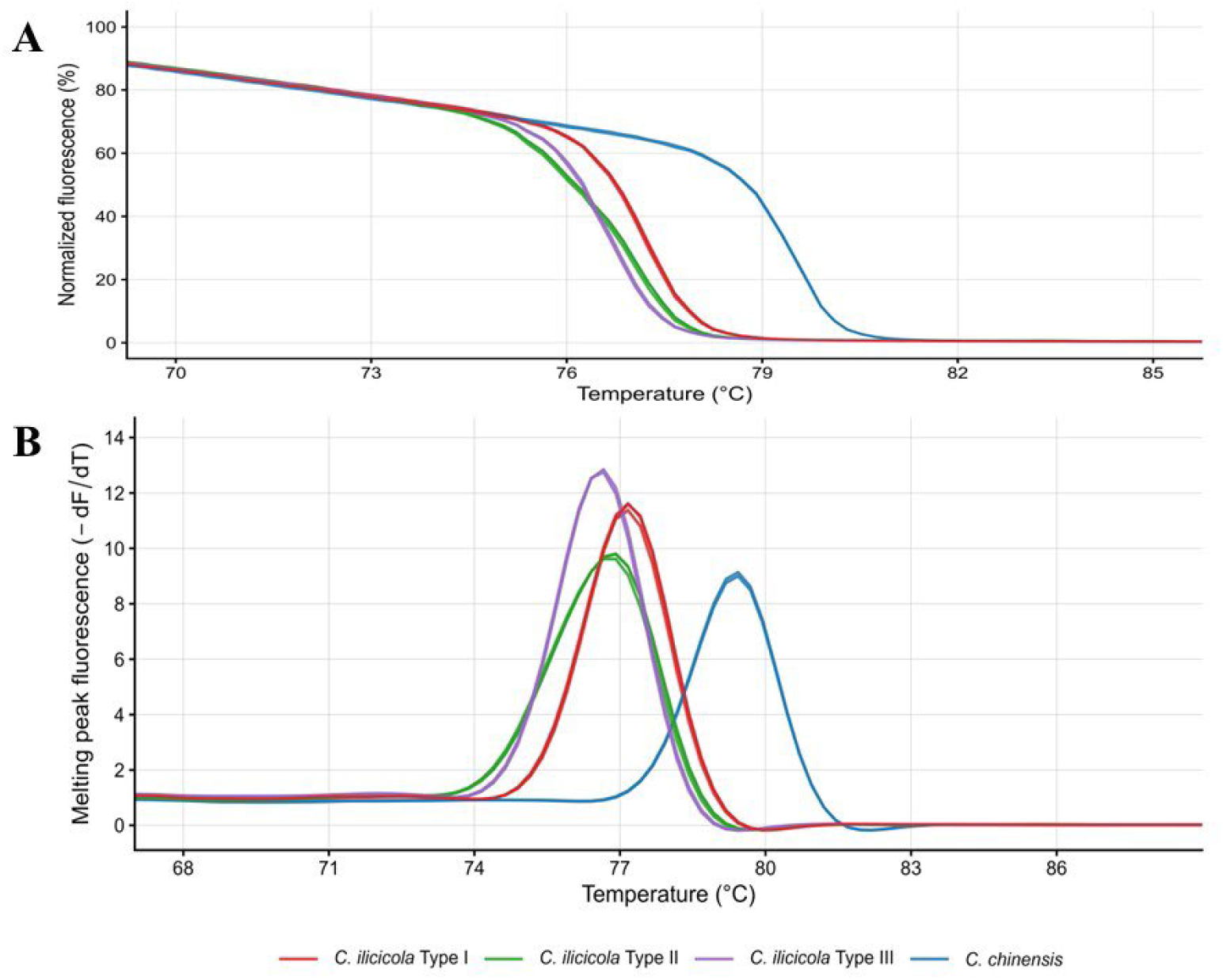
HRM profiles of *C. ilicicola* isolates and *C. chinensis*. (A) Normalized melting curves and (B) derivative melting peaks illustrating three characteristic HRM profile types among *C. ilicicola* isolates, along with a distinct melting profile observed for the non-target species *C. chinensis*. *C. ilicicola* Type I isolates consistently produced melting peaks near 77 °C, whereas Type II and Type III isolates produced peaks near 76 °C but were distinguishable by differences in peak shape and height. In contrast, *C. chinensis* yielded a clearly separated melting peak near 79 °C, indicating strong assay specificity.

### HRM assay differentiates *C. ilicicola* isolates

HRM analysis revealed clear variation among *C. ilicicola* isolates, enabling discrimination among three haplotypes present within the 73-bp TEF-1α amplicon, with three distinct melting-profile groups detected (Figure 2). The most common group, consisting of isolates from Illinois and surrounding states, produced a sharp, uniform melting peak centered near 77°C (Type I). A second group melted near 76°C but generated broader, flattened, and more symmetrical melting profiles (Type II). A third group also exhibited slightly lower melting temperatures (∼76 °C) but produced sharp and uniform peaks (Type III). Isolates producing broader peaks amplified later and showed higher Cp values, a pattern consistent with the presence of a single-nucleotide deletion in the reverse-primer binding region, which may reduce primer-binding efficiency. In contrast, isolates with sharper peaks amplified earlier and produced stronger fluorescence signals, reflecting the absence of primer-binding mismatches. The observed melting-profile differences corresponded directly with sequence variation in the TEF-1α target region, including the G→A single-nucleotide polymorphism and the one-base deletion detected in a subset of isolates (Figure 1). Collectively, the HRM assay successfully resolved all three *C. ilicicola* haplotypes, demonstrating its capacity to detect isolate-specific variations.

## HRM and PCR assay from infected plant tissue

A total of 56 samples were collected from soybean fields in Illinois based on field observations suggestive of red crown rot (RCR). Of these, 36 displayed clear RCR symptoms, 11 showed slight discoloration in the crown region, and 9 asymptomatic samples were also included. Consistent with the observed symptoms, HRM analysis detected *C. ilicicola* in all 36 samples that displayed clear RCR symptoms. HRM-positive samples produced melting profiles similar to those of the *C. ilicicola* genomic DNA positive control and were clearly distinguishable from negative controls (Figure 3). The remaining 20 samples were not classified as *C. ilicicola* by HRM, in agreement with visual symptom assessments. These samples exhibited non-matching melting curves, occasionally displaying broader or shifted peaks, as well as weak or late amplification approaching 40 cycles. Because these melting patterns did not align with the *C. ilicicola* positive-control profile, they were interpreted as negative. To further validate HRM-based detection, a subset of 20 field samples representing both HRM-positive and HRM-negative outcomes was analyzed using conventional PCR targeting the IGS region, followed by gel electrophoresis and Sanger sequencing (Supplementary Figure 4). Samples classified as positive by HRM produced amplicons of the expected size, and sequencing confirmed their identity as *C. ilicicola*. Samples classified as negative by HRM did not yield *C. ilicicola* sequences. Collectively, the strong agreement among HRM results, field symptomology, conventional PCR, and sequencing demonstrates that the HRM assay provides reliable detection of *C. ilicicola* directly from infected soybean samples, eliminating the need for pathogen isolation or culturing.

**Figure 3.**
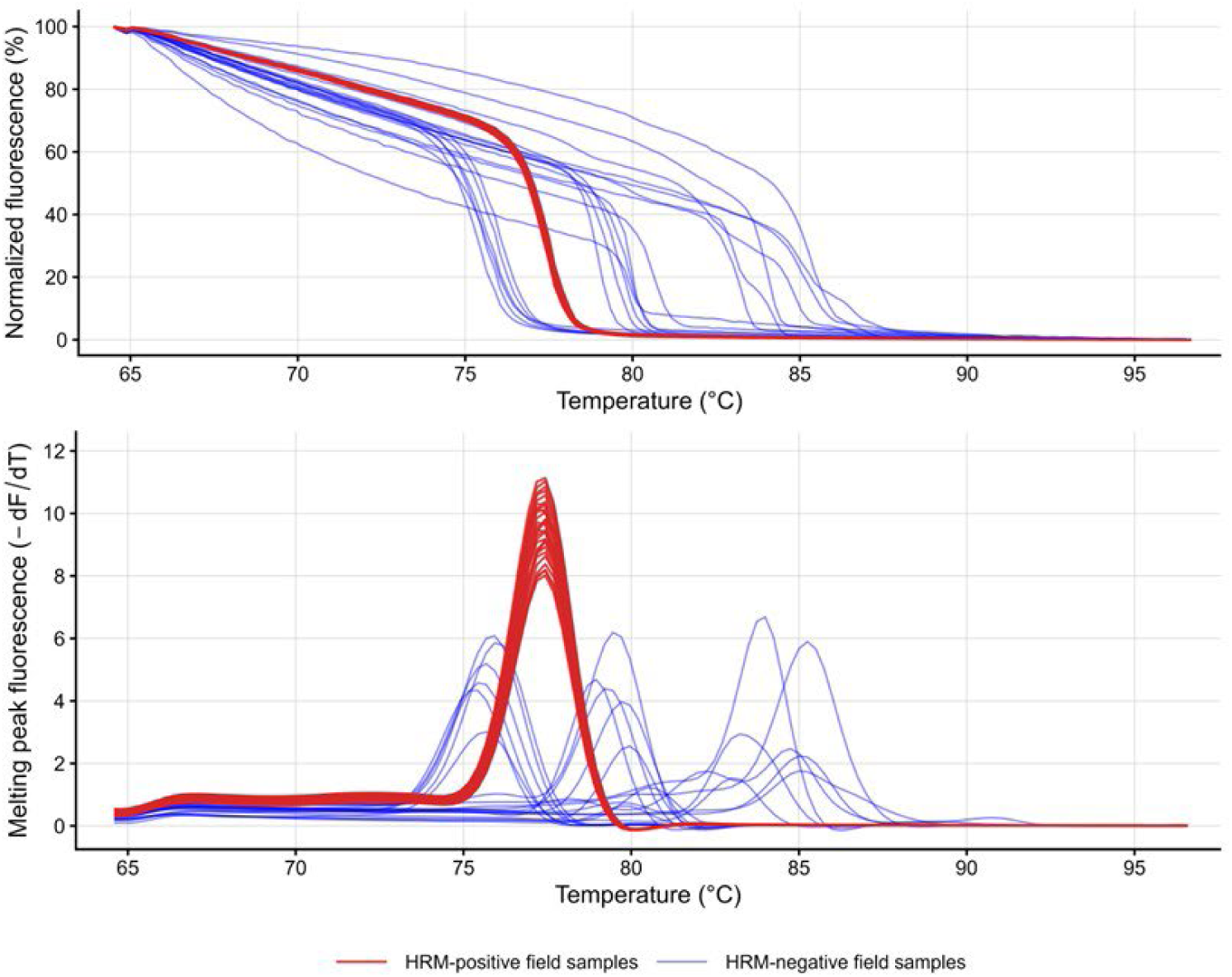
HRM analysis of field samples tested for *C. ilicicola*. Normalized melting curves and derivative melting peaks from HRM-positive field samples (red) closely matched the characteristic *C. ilicicola* positive-control profile, producing reproducible melting peaks centered near 77 °C. In contrast, DNA from asymptomatic soybean plants (blue) exhibited late-cycle amplification and produced shifted, broader, weaker, or otherwise inconsistent melting profiles, and were therefore not classified as *C. ilicicola*. These results demonstrate that the HRM assay reliably detects *C. ilicicola* directly from infected soybean tissue while distinguishing non-target variable amplifications associated with non-symptomatic plants.

## Discussion

*C. ilicicola*, the causal agent of RCR, is an emerging threat in the U.S. Midwest region, and accurate identification is critical because field symptoms often resemble those caused by other soilborne diseases. We developed a high-resolution melting (HRM) assay that accurately distinguishes *C. ilicicola* from other fungi and oomycetes as well as intraspecific variation among the isolates based on characteristic melting profiles. This enables rapid pathogen detection directly from infected soybean tissues and providing an efficient alternative to traditional culture-based diagnostic methods.

For genotyping assays, selecting a genomic region that is conserved across diverse isolates ensures consistent amplification, while sequence variation within the amplified fragment enables isolate-level discrimination. HRM analysis is well suited for this purpose because even a single nucleotide change can generate a distinct melting profile. As a closed-tube, cost-effective, and highly sensitive method, HRM provides a practical alternative to sequencing or probe-based qPCR assays for large-scale genotyping efforts. Multicopy ribosomal targets can improve analytical sensitivity because rRNA genes occur in higher copy numbers in genomes (Black et al. 2013). However, when ribosomal regions do not provide sufficient resolution among closely related fungi, protein-coding genes such as TEF-1α can provide useful sequence variation for species-level discrimination (Roger et al. 1999; Yörük and Yli-Mattila 2024). In silico analysis of TEF-1α sequences from soybean root-associated fungi and oomycetes, *Calonectria* spp., and *C. ilicicola* isolates confirmed that this locus provides an optimal balance of conservation and variability. In our assay, HRM differentiation among *C. ilicicola* isolates corresponded directly to sequence differences in the TEF-1α target region, including a SNP within the amplicon and a one-nucleotide deletion at the reverse primer-binding site. These variants produced distinct melting behaviors, demonstrating the ability of HRM not only to detect *C. ilicicola* but also to resolve naturally occurring genetic variation among isolates.

Quantitative PCR methods can generate false positives due to non-specific binding, primer-dimer formation, or interference with the PCR reaction (Navarro et al. 2015). These limitations can be mitigated through careful assay design, appropriate controls, and the use of optimized buffers and saturating dyes (Bustin et al. 2025). Effective implementation also requires recognition that HRM interpretation is context-dependent, with thresholds and criteria for defining positive results varying according to assay design, target organism, and sample type (Słomka et al. 2017). Small differences in DNA quality or reaction setup can influence amplification efficiency, causing shifts in Cp values that ultimately affect melting-curve profiles. Because HRM clustering is highly sensitive to these fluctuations, consistent sample preparation and reaction conditions are essential for reliable interpretation. Published HRM studies reflect this variability, using 30 cycles for *Leptosphaeria maculans* AvrLm4-7 genotyping (Carpezat et al. 2014), 35 cycles for SSR-HRM analysis of *Venturia inaequalis* (Chatzidimopoulos et al. 2019), detection of multiple fungal species that cause crown rot diseases in strawberry (Wang et al. 2021), *Fusarium graminearum* chemotype differentiation (Singh et al. 2024), and 40 cycles for differentiation of *Alternaria* species causing brown spot disease in citrus (Garganese et al. 2018). Collectively, these studies demonstrate that cycle number alone cannot define HRM positivity; amplification timing must be interpreted together with melting-curve characteristics to ensure accurate detection. Consistent with this principle, our study evaluated specificity using the full melting profile rather than relying on a single metric. For pure DNA samples, we applied a stringent 30-cycle cutoff, and our results showed Cp values of ∼18 cycles for *C. ilicicola* DNA (20 ng/µL). In contrast, plant tissues in our study that were infected with *C. ilicicola* had similar melting profiles with Cp at 22-30 cycles, indicating higher cycle thresholds may be necessary for infected plant tissues where pathogen DNA is present at lower abundance relative to host DNA.

The most effective molecular detection methods for plant pathogens are those capable of detecting or genotyping the pathogen directly from infected plant tissue, eliminating the need to isolate and culture the organism. This greatly increases throughput and reduces the time and resources required for reliable identification. However, direct detection from host tissues presents challenges because plant genomic DNA is abundant and plant tissues contain molecules that can inhibit downstream molecular assays (Malvick and Grunden 2005; Wei et al. 2008; Yang et al. 2016). Despite these complexities, several studies have demonstrated the effectiveness of HRM assays for direct pathogen detection from host tissues, including identification of pathogens causing crown rot and Pestalotia leaf spot in strawberry (Rebello et al. 2023; Wang et al. 2021), detection of *Monilinia* spp. responsible for brown rot in peach (Papavasileiou et al. 2016), and chemotype characterization of the *Fusarium* head blight pathogen in wheat (Singh et al. 2024). Applying this approach to soybean field samples exhibiting RCR-like symptoms, our HRM assay reliably detected *C. ilicicola* in patterns consistent with disease symptoms. In agreement with previous reports of low *C. ilicicola* genetic diversity in Illinois and nearby states (Clough et al. 2026), only Type I isolates were identified in field samples. Field-derived DNA generally produced slightly higher Cp values than pure-culture DNA, which is expected because pathogen abundance is lower in plant tissues and plant-derived inhibitors can reduce amplification efficiency (Rački et al. 2014). In our study, Cp values from infected soybean tissues ranged from 22 to 30 across 36 samples. Similar variability in pathogen DNA levels has been documented in soybean sudden death syndrome (SDS), where differences in disease severity, tissue type, and inhibitor content influence amplification timing (Gao et al. 2004).

Nevertheless, samples from plants displaying RCR symptoms consistently produced melting profiles identical to reference *C. ilicicola* DNA and were thus classified as positive, whereas samples yielding weak, shifted, or atypical melt curves were designated as negative. HRM-based detection results were further validated using conventional PCR targeting the IGS region, followed by Sanger sequencing to confirm amplicon identity. Together, these results demonstrate that the HRM assay provides a robust, rapid, closed-tube approach for detecting *C. ilicicola* directly from soybean tissue. Beyond diagnostic accuracy, the consistent HRM profiles across isolates underscore the assay’s promise for broader molecular surveillance. As *C. ilicicola* continues to expand geographically, routine monitoring will be essential for tracking changes in pathogen prevalence and genotype distribution. Although the assay performed reliably across diverse sample types, HRM analyses can be affected by DNA quality, dye chemistry, and instrument-specific melting resolution (Druml and Cichna-Markl 2014). Future work should include cross-laboratory and cross-platform validation, similar to the multi-laboratory qPCR comparison conducted for SDS (Kandel et al. 2015), as well as expansion of the isolate panel to encompass additional geographic regions. These efforts would enable finer-scale assessment of intraspecific diversity and further refine assay performance, ultimately supporting long-term monitoring and management of RCR.

In conclusion, we developed and validated an HRM assay targeting the TEF-1α region for rapid, cost-effective, and accurate detection of *C. ilicicola*. The assay showed strong analytical performance, reliably distinguishing *C. ilicicola* from non-target species and successfully detecting the pathogen directly from infected soybean tissue. Additionally, sequence variation within the targeted region produced distinct HRM profiles among isolates, enabling detection of intraspecific variation and demonstrating the assay’s potential for future applications in exploring genetic diversity with biological and evolutionary significance.

## ACKNOWLEDGEMENTS

We thank Theresa Herman (USDA-ARS, Urbana, IL) for maintaining the *C. ilicicola* isolates and Nancy McCoppin (USDA-ARS, Urbana, IL) for managing laboratory supplies and assisting with buffer preparation. We also thank Dr. Lisa Keith (USDA-ARS, Hilo, HI), Dr. Santiago Mideros (University of Illinois, Urbana–Champaign, IL) and Dr. Ahmad Fakhoury (Southern Illinois University, Carbondale, IL), Dr. Nina Shishkoff and Dr. Douglas Luster (USDA-ARS, Fort Detrick, MD) for providing a subset of fungal isolates and DNA used in this study, as well as David Neece (USDA-ARS, Urbana, IL) for providing soybean genomic DNA used in the study. Mention of trade names or commercial products in this article is solely for the purpose of providing specific information and does not imply recommendation or endorsement by the U.S. Department of Agriculture (USDA). USDA is an equal opportunity provider and employer.

## Funding

Support was provided by the U.S. Department of Agriculture–Agricultural Research Service (USDA-ARS) CRIS project 5012-22000-023-000-D, and Pioneer Fellowship, James B. Sinclair Fellowship, and Johnny W. Pendleton Fellowship to K. R. Pandey from the Department of Crop Sciences at University of Illinois Urbana- Champaign.

## Notes

### Competing Interest Statement

The authors have declared no competing interest.

