## Supplementary Figure for "High-Resolution Melting (HRM) Assay for Molecular Detection of *Calonectria ilicicola* in Soybean"

### Supplementary Figure Legends

**Supplementary Figure S1. Representative soybean root samples showing symptomatic and asymptomatic plants included in this study for HRM analysis.** The figure illustrates the range of root conditions used for assay evaluation, including plants exhibiting characteristic RCR symptoms and asymptomatic plants collected as negative controls. These representative samples demonstrate the diversity of field material incorporated into HRM-based detection of *C. ilicicola*.

**Supplementary Figure S2. Standard curve of *C. ilicicola* DNA generated using the HRM primer pair.** Mean Cp values from four technical replicates were plotted against the $\log_{10}$ DNA concentration (ng/µL). The resulting regression equation ($y = -3.473x + 21.1$), coefficient of determination ($R^2 = 0.9992$), and amplification efficiency (94.05%) demonstrate strong linearity and reliable quantitative performance across the tested concentration range.

**Supplementary Figure S3. Amplification curves from the HRM assay using DNA extracted from pure cultures included in the specificity panel.** Amplification profiles are shown for all fungal species and *C. ilicicola* isolates tested. *C. ilicicola* Type I and Type III isolates exhibited early amplification, whereas Type II isolates amplified later relative to Type I/III. *C. chinensis* produced intermediate amplification, and non-target species exhibited late, weak, or inconsistent amplification curves. Amplification occurring after 30 cycles was considered non-specific under these assay conditions.

**Supplementary Figure S4.** Agarose gel electrophoresis of PCR amplification using IGS-specific primers. The gel displays grouped samples as indicated above the lanes: soybean samples infected with RCR (labeled 1–7), asymptomatic soybean samples lacking RCR (labeled 8–12), and soybean cultivars included for comparison (labeled 13–16). A water (no-template) control (W) and a positive control containing *C. ilicicola* DNA (P) are included. DNA ladders are shown on the left and right sides of the gel, with the 500-bp band marked for reference.

**Supplementary Figure S1.**

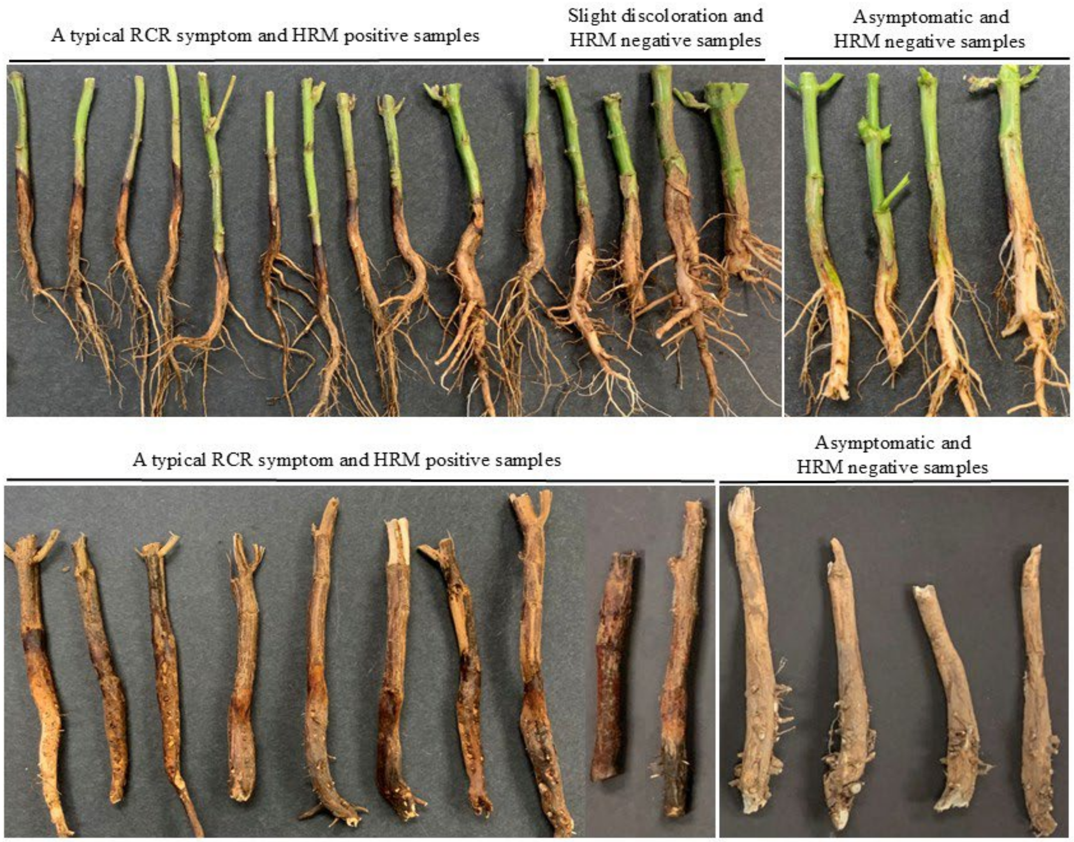

**Supplementary Figure S2.**

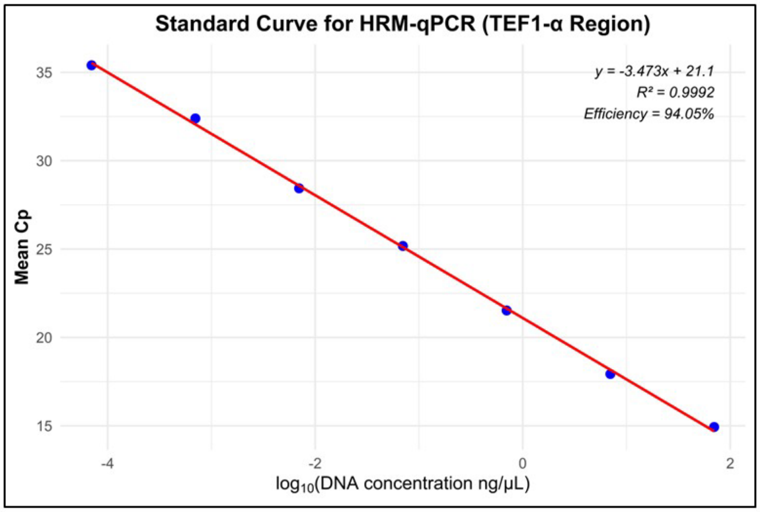

**Supplementary Figure S3.**

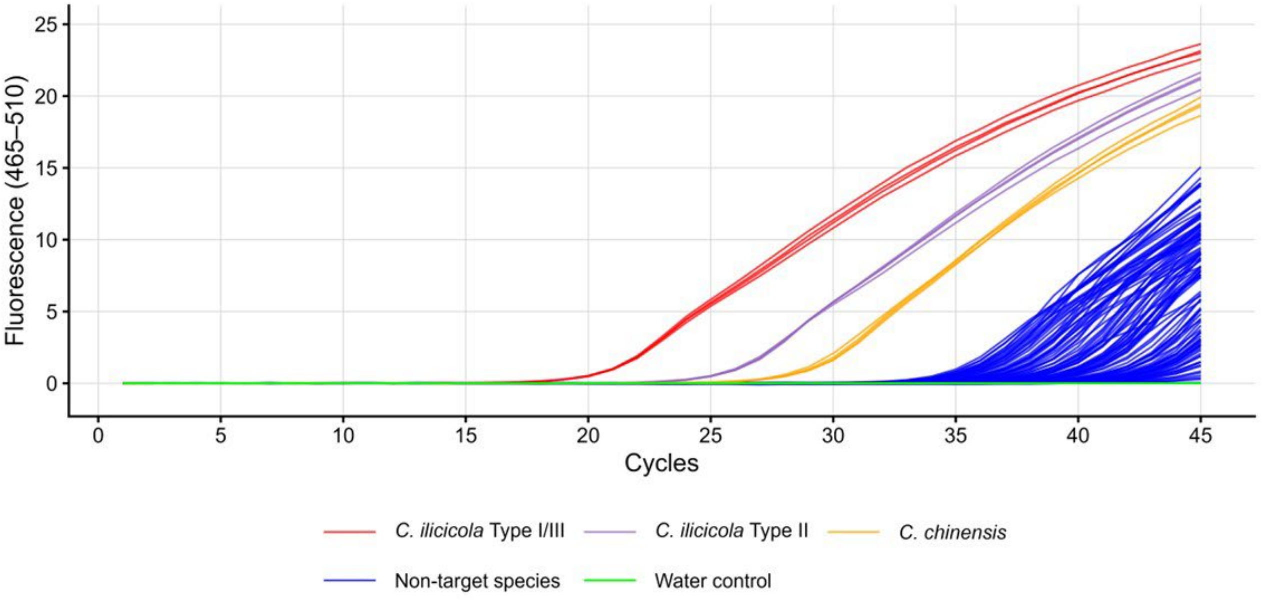

**Supplementary Figure S4.**

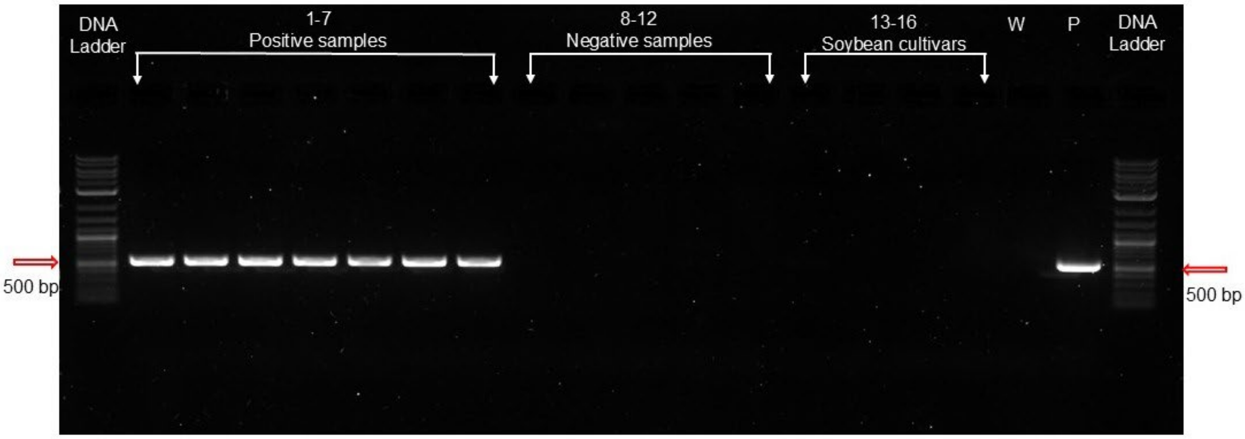
